# Emergence of Small Colony Variants (SCVs) is an adaptive strategy used by *Pseudomonas aeruginosa* to palliate O_2_ limitations

**DOI:** 10.1101/2022.07.20.500912

**Authors:** Alison Besse, Marie-Christine Groleau, Eric Déziel

## Abstract

The ability to generate a subpopulation of Small Colony Variants (SCVs) is a conserved feature of *Pseudomonas aeruginosa* and could represent a key adaptive strategy to colonize and persist in multiple niches. However, very little is known about the role of the SCV phenotype, the conditions that promote its emergence and its possible involvement in an adaptive strategy. In the present work, we investigated the *in vitro* selective conditions promoting the emergence of SCVs from the prototypical strain PA14, which readily forms SCVs in non-agitated standing cultures. We found that O_2_ limitations, which cause a redox imbalance, is the main factor selecting for the SCV phenotype, which promotes survival of the population *via* formation of a biofilm at the air-liquid interface to access the electron acceptor. When this selective pressure is relieved by aeration or supplementation of an alternative electron acceptor, SCVs are barely detectable. We also observed that SCV emergence contributes to redox rebalancing, suggesting that it is involved in an adaptive strategy. We conclude that selection for the SCV phenotype is an adaptive solution adopted by *P. aeruginosa* to access poorly available O_2_.

**IMPORTANCE:** The bacterium *P. aeruginosa* is an opportunistic pathogen that thrives in many environments. It poses a significant health concern, notably because it is a causative agent of nosocomial infections and the most prevalent pathogen found in the lungs of people with cystic fibrosis. In infected hosts, its persistence is often related to the emergence of an alternative phenotype known as Small Colony Variant (SCV). Identification of conditions selecting for the SCV phenotype contributes to knowledge regarding adaptive mechanisms exploited by *P. aeruginosa* to survive in multiple niches and persist during infections. Hindering this adaptation strategy could help control *P. aeruginosa* persistent infections.

## INTRODUCTION

The bacterium *Pseudomonas aeruginosa* thrives in diverse environments, such as aquatic habitats, soil, food, and even built environments, such as hospital premise plumbing systems (1–5). In addition to environmental habitats, *P. aeruginosa* strains are frequently isolated from clinical samples. It is one of the most frequent causative agents of nosocomial infections and a major cause of infections among immunocompromised individuals, especially in people with cystic fibrosis (CF) (6–8). *P. aeruginosa* is also capable of infecting non-mammalian hosts, such as plants and insects (9–11). To persist in these various environments, *P. aeruginosa* has to face hostile conditions such as poor nutrient availability, O_2_ limitation, and desiccation, along with host immunity and antimicrobial treatments in clinical context.

The successful adaptation of *P. aeruginosa* to multiple niches is attributed to its high genomic and metabolic versatility (12). For instance, the flexible metabolism of *P. aeruginosa* allows it to use different final electron acceptors to produce energy. Although energy generation is based mainly on oxidative substrate catabolism, *P. aeruginosa* is able to grow under anaerobic conditions with alternative electron acceptors, such as nitrate, or to survive by fermentation of arginine or pyruvate (13–15). Activation of the denitrification pathway is regulated by ANR, a direct O_2_ sensor that regulates anaerobic gene expression (16). When *P. aeruginosa* undergoes electron acceptor limitations during *in vitro* biofilm growth, it develops different strategies to palliate low-O_2_ conditions. These strategies include the production of small redox molecules, called phenazines, which act as electron carriers shuttling electrons from intracellular metabolism to distant extracellular oxidants (17, 18). Phenazines behave as alternative electron acceptor, and their production facilitate intracellular redox balancing by oxidizing redox state (19). *P. aeruginosa* can also cope with O_2_ limitation by changing the overall structure of a biofilm to increase the surface exposed to O_2_ (20, 21); this is visible as wrinkles on the surface of colony biofilms (20).

The virulence and persistence of *P. aeruginosa* in various environments are also mediated by phenotypic and genotypic diversification, generating distinct sub-populations. Phenotypic adaptations have been especially studied in the clinical infection context, notably CF (22, 23). *P. aeruginosa* isolates from CF lungs display distinct phenotypic variations, such as Small Colony Variants (SCVs) (24, 25). Emergence of SCVs is a frequently observed feature of *P. aeruginosa* isolates from infected patients, especially CF individuals in which SCV emergence is correlated with persistent infection (25–30). SCVs are not restricted to clinical context and the ability to adopt a SCV phenotype is widely distributed among *P. aeruginosa* strains. Strains isolated from various non-clinical environments such as soil, food, hospital water systems can also form SCVs (31, 32). Since the ability of *P. aeruginosa* to form SCVs is a conserved feature, this could represent a key adaptation strategy to colonize and persist in multiple niches. Furthermore, SCVs can be readily selected under *in vitro* laboratory conditions (31–33). Thus, selection of the SCV phenotype could be exploited by *P. aeruginosa* to respond to various stressful conditions.

However, knowledge is missing regarding the specific conditions that promote SCVs emergence, or the benefit of their selection in a possible adaptive strategy. In this work, we investigated the selective pressure promoting SCV emergence in *in vitro* experiments. We chose the prototypical strain PA14, which we found readily forms SCVs in non-agitated standing culture conditions (32) to investigate the effect of O_2_ limitations on SCV emergence and survival by promoting the formation of a biofilm at the air-liquid interface.

## RESULTS

### Pellicle formation is linked to SCV incidence

SCVs of *P. aeruginosa* are obtained under culture conditions promoting biofilm formation (25, 31, 33). Accordingly, we do not recover SCVs from standard agitated cultures of the prototypical clinical strain *P. aeruginosa* PA14 even after 48 h (Fig. 1A) and there is no formation of a pellicle at the air-liquid interface (Fig 1B). However, when PA14 is grown as a standing culture for 48 h, SCVs emerge spontaneously (Fig 1A). Since a surface pellicle is noticeable in standing cultures (Fig 1B), these results indicate that SCVs emerge under conditions promoting biofilm formation.

**Fig. 1.**
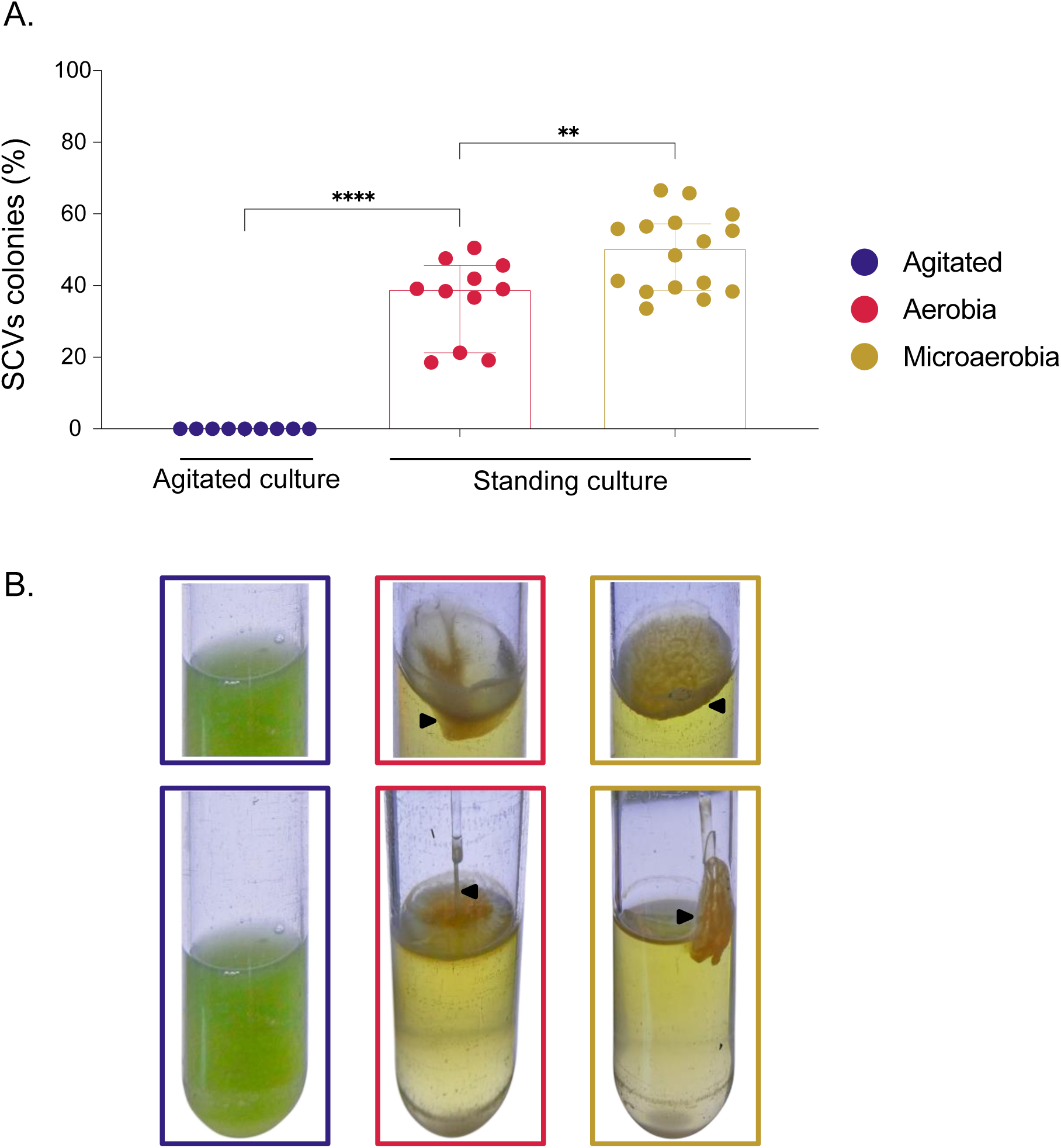
SCVs rate and growth pattern of cultures depending on O_2_ availability. *Pseudomonas aeruginosa* PA14 was cultivated in agitated, in static aerobic or in static microaerobic conditions in TSB. (A) After 16h (agitated) or 48h (static) of incubation, ratios of SCVs colonies were counted. Results are expressed in % of total population. Each dot represents counts for one independent culture; (B) Growth pattern of cultures incubated in agitated (purple), static aerobic (pink) and static microaerobic (yellow) conditions. Black arrows indicate pellicle formed at the air-liquid surface. Pellicles were removed with a Pasteur pipette tips to gauge their cohesive properties. Note that counts and pellicles were also assessed at 48h for the agitated culture with similar results. The statistical significance of the results was calculated by a non-parametric Mann-Whitney test, ****, P value ≤ 0.0001; **, P value ≤ 0.01.

Next, we asked whether biofilm-promoting conditions are exclusively responsible for the emergence of SCVs or if an additional environmental pressure could be involved. A key difference between agitated and standing cultures is the O_2_ distribution within the liquid. Thus, we investigated the role of O_2_ in this phenomenon by cultivating *P. aeruginosa* PA14 under microaerobic conditions; strikingly, an even more hydrophobic and robust pellicle developed at the surface, as shown by its sticky properties when removing it with a plastic tip (Fig 1B). Indeed, 39 % of the total population of a 48 h aerobic standing culture were SCVs (Fig 1A), while when the environmental O_2_ concentration was reduced, i.e. under microaerobic conditions, SCVs reached 50% of the total population (Fig 1A). Since microaerobic conditions promote a higher SCV proportion (Fig 1A) and that SCVs are highly hydrophobic, this result confirms that pellicle formation is directly correlated with SCVs incidence.

### Rapid depletion of oxygen in static cultures selects for SCV emergence

To further verify if there is a link between O_2_ availability and the incidence of SCVs in the culture, we added resazurin dye to 5 mL standing cultures. The dye turned pink as early as 6 h after inoculation, indicating that O_2_ was rapidly consumed by the bacteria (Fig 2A). After 48 h of incubation, O_2_ was mostly depleted, as shown by the colorless (i.e. reduced) resazurin in most of the culture and it was oxidized at the surface only (top pink layer). This indicated that O_2_ was rapidly consumed by the cells, and that only surface cells could have access to significant O_2_ (Fig 2A). Interestingly, SCVs were forming a significant portion of the whole population as soon as 24 h after the beginning of the incubation, concomitantly with the observation of a significant turbidity and prior to complete O_2_ depletion (Figs. 2A and 2B). The % of SCVs within the total population reached its maximum after 48 h of incubation, when the O_2_ was almost completely depleted (Fig 2A and B). Since O_2_ diffusion was limited to the superficial part of the PA14 standing culture after 48 h, we sampled 1 ml fractions from the top to the bottom of the standing culture and quantified SCVs (Fig 2C). SCVs represented ~30% of total population in the three upper fractions, where the resazurin was still oxidized, but this % dropped significantly to ~15% in the deeper fractions 4-8 (Fig 2C). Thus, SCVs were predominantly found at the surface, where O_2_ was still detectable and SCVs incidence was drastically reduced in parts of the culture where O_2_ was depleted (Fig 2C).

**Fig. 2.**
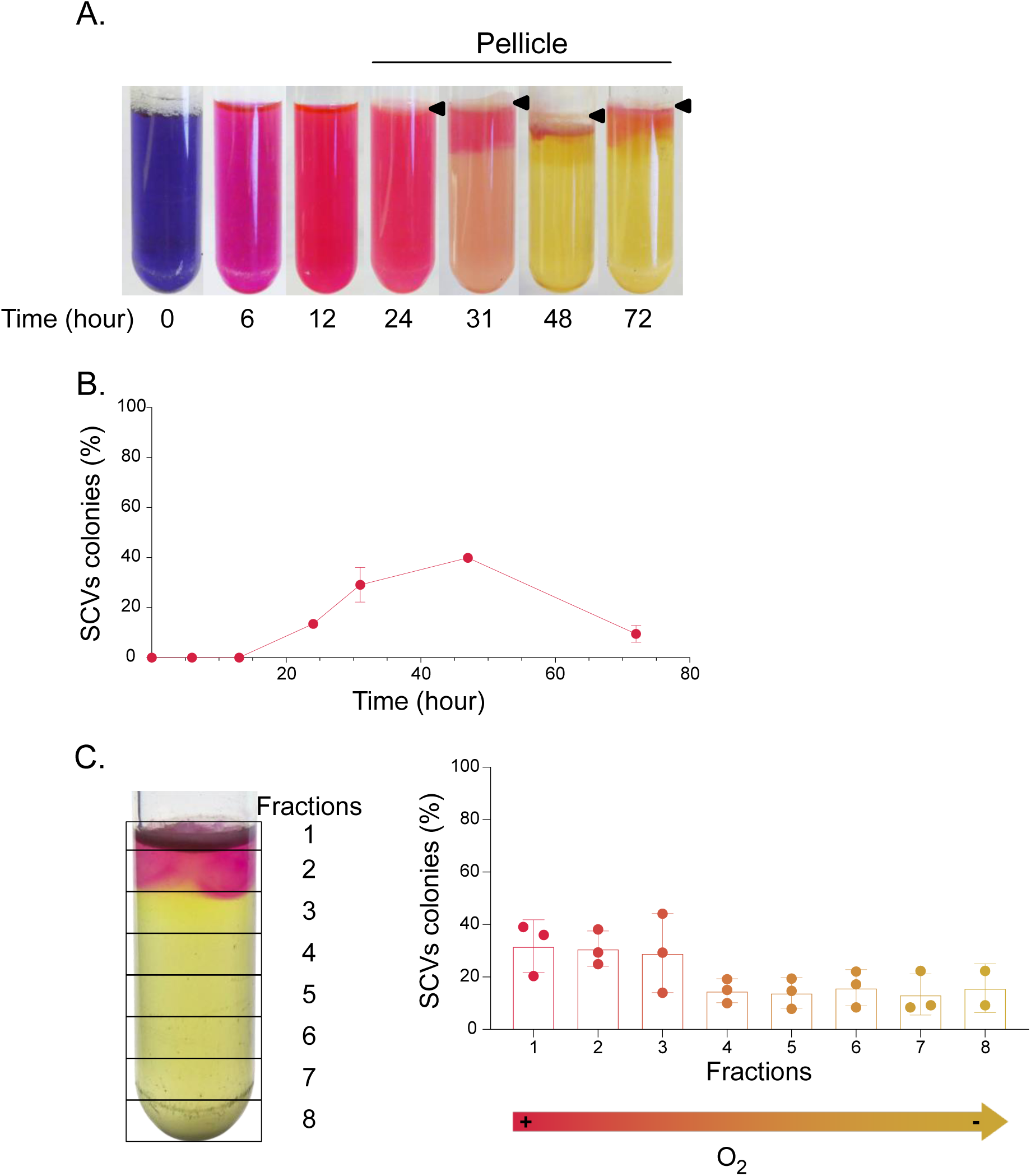
SCVs emergence and distribution in standing cultures depending on O_2_ depletion over time. PA14 was inoculated in TSB with 100 mg/ml of resazurin and incubated in aerobic static conditions. (A) After 0, 6, 12, 24, 31, 48 and 72 h, resazurin reduction was observed to assess O_2_ depletion within culture. Black arrows point to the pellicle formed at the air-liquid surface. (B) After 0, 6, 12, 24, 31, 48 and 72 h, cultures were sampled and SCVs colonies were counted and expressed in % of total population. (C) After 50 h of incubation, 1 ml fractions form the top to the bottom of the culture were sampled, spread onto TSA plates and SCVs were quantified (% of total population).

Since SCVs promote efficient biofilm growth, we hypothesized that SCVs are functionally selected because they readily form a pellicle at the air-liquid interface, allowing rapid adaptation of the population to electron acceptor limitation by providing access that is more direct to atmospheric O_2_. Supporting this hypothesis, *P. aeruginosa* PA14 grows mainly as a pellicle in O_2_-limiting conditions, with almost no turbidity visible under the pellicle (e.g. Fig 1B).

Finally, a lag phase of ~20 h was observed prior to efficient growth in standing culture (Fig. 3C). Accordingly, it is also the timing observed before the first detection of SCVs and visible occurrence of a pellicle in standing cultures (Fig. 2B), suggesting that during the lag phase observed, an initially minor SCV subpopulation is strongly positively selected to overcome survival pressure caused by O_2_ restriction. All together, these results indicate that restricted access to O_2_ represents a strong selective pressure to switch to the SCV phenotype in *P. aeruginosa* cultures.

**Fig. 3.**
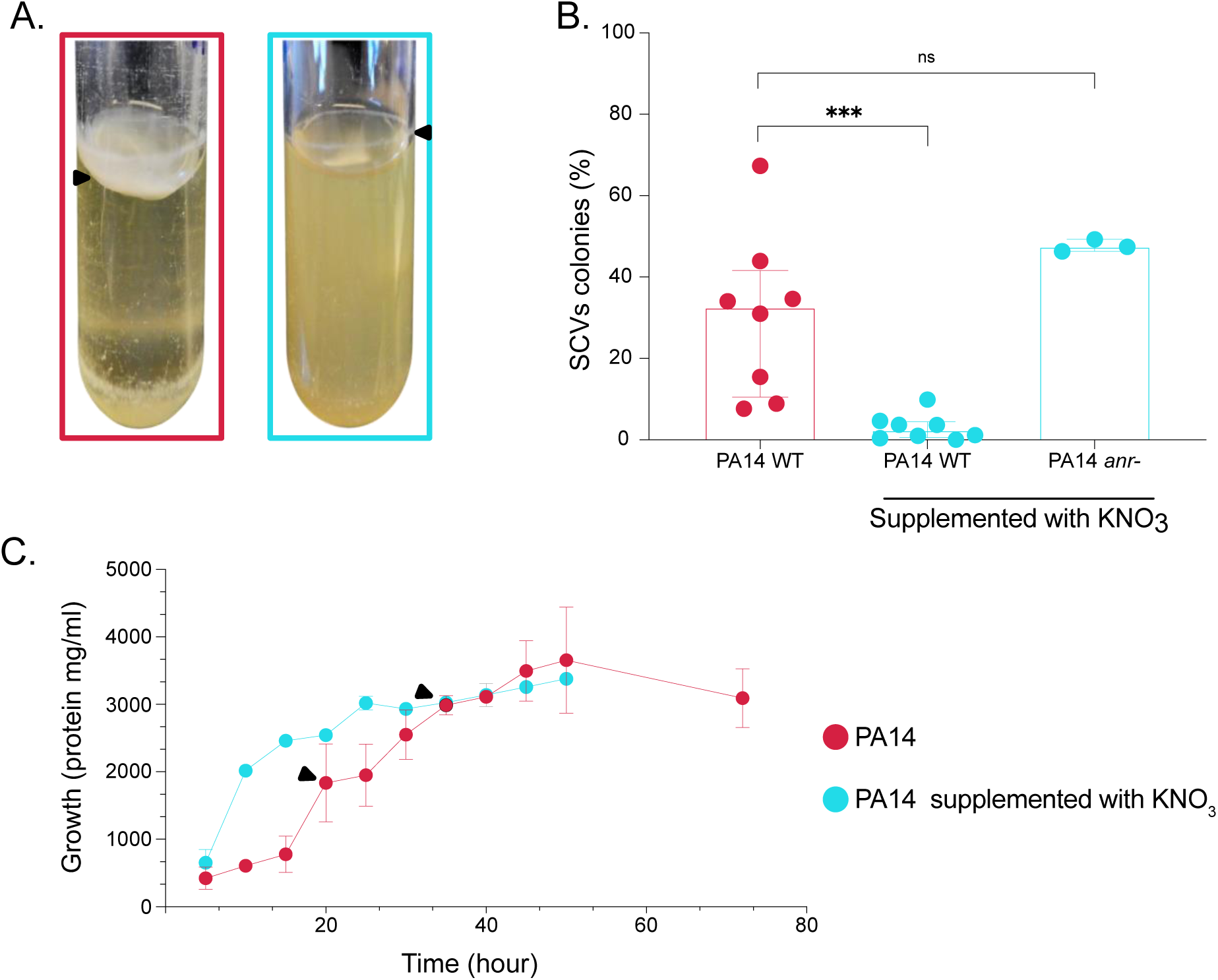
Growth pattern and SCVs emergence upon supplementation with nitrate. PA14 was inoculated in TSB or TSB supplemented with 50 mM KNO_3_. (A) Growth pattern and pellicle formation in PA14 culture not supplemented (pink) or KNO_3_ supplemented (turquoise blue). Black arrows indicate pellicle or residual pellicle formed at the surface air-liquid. (B) PA14 WT and PA14 *anr-* were inoculated in TSB or TSB supplemented with KNO_3_. After 48 hours of incubation, cultures were spread onto TSA plates and % of SCVs were quantified. Results are expressed in % of SCVs in total population. The statistical significance of the results calculated by a non-parametric Mann-Whitney test, ***, P ≤ 0.001; ns, not significant. (C) Growth of PA14 in TSB and TSB-KNO_3_ over time. Black arrows indicate first detection of a pellicle at the air-liquid interface.

### Presence of an homogeneously available alternative electron acceptor relieves the selective pressure to switch to SCVs

We next wanted to determine if the selection, caused by heterogenic O_2_ distribution in PA14 standing culture, was indeed responsible for SCV emergence, but also for pellicle formation. This was verified by adding the alternative electron acceptor nitrate (as KNO_3_) to standing cultures, so that O_2_ was no longer the primary electron acceptor. Supporting our hypothesis, no clear pellicle was observed in presence of KNO_3_, only a residual trace of a thin, fragile pellicle (Fig. 3A). This residual pellicle was nothing like the cohesive thick pellicle observed without presence of nitrate (Fig. 3A). Also, with KNO_3_ supplementation, the culture medium was turbid, indicating that the growth was homogeneous within the culture and no longer mainly localised at the surface as observed without addition of KNO_3_ (Fig. 3A).

Strikingly, SCVs were barely detectable in standing cultures with KNO_3_ (Fig. 3B), indicating that presence of an homogeneously available electron acceptor reduced SCVs emergence. Anr is the main O_2_ regulator of *P. aeruginosa* and is activated by low O_2_ conditions to control expression of the denitrification genes, allowing the use of nitrate as electron acceptor. Thus, *P. aeruginosa anr* mutants are no longer able to use nitrate as an O_2_ alternative electron acceptor. Accordingly, a *P. aeruginosa* PA14 *anr*^_*-*_^ mutant produced as much SCVs as the WT when KNO_3_ was present (Fig. 3B). All together, these results indicate that when O_2_ limitation is overcome, the pressure to grow at the surface as a pellicle and thus switch to SCV phenotype is removed, consistent with the absence of SCVs in well-agitated cultures (Fig. 1A). Limitations in an homogeneously available electron acceptor, rather than specific O_2_ limitation, is the selective pressure resulting in SCVs emergence and pellicle formation under our conditions.

### SCVs emergence is promoted in the absence of an effective electron shuttle to access O_2_

As previously mentioned, in biofilms and biofilm-like microbial communities, access to O_2_ is often restricted and generally localized at the surface of the growing structure. To overcome this limitation, *P. aeruginosa* can channel electrons from intracellular metabolism to distant extracellular oxidants, through electron carriers such as phenazines (34). Interestingly, growth in standing cultures of a ∆∆*phzC1C2* mutant, unable to produce phenazines, results a higher SCVs proportions (Fig. 4A). The same result was observed for a *phzM* mutant, enable to convert phenazine-1-carboxylic acid (PCA) to pyocyanin, a blue pigmented phenazine. Complementation of these *phz* mutants with exogenously supplemented pyocyanin inhibited the emergence for SCVs, indicating that specific absence of pyocyanin production by these mutants promotes emergence of SCVs (Fig. 4, A). Accordingly, increasing concentrations of added pyocyanin to WT standing cultures negatively correlated with the emergence of SCVs (Fig. 4B), supporting a central role for this phenazine in accessing distantly available electron acceptor.

**Fig. 4.**
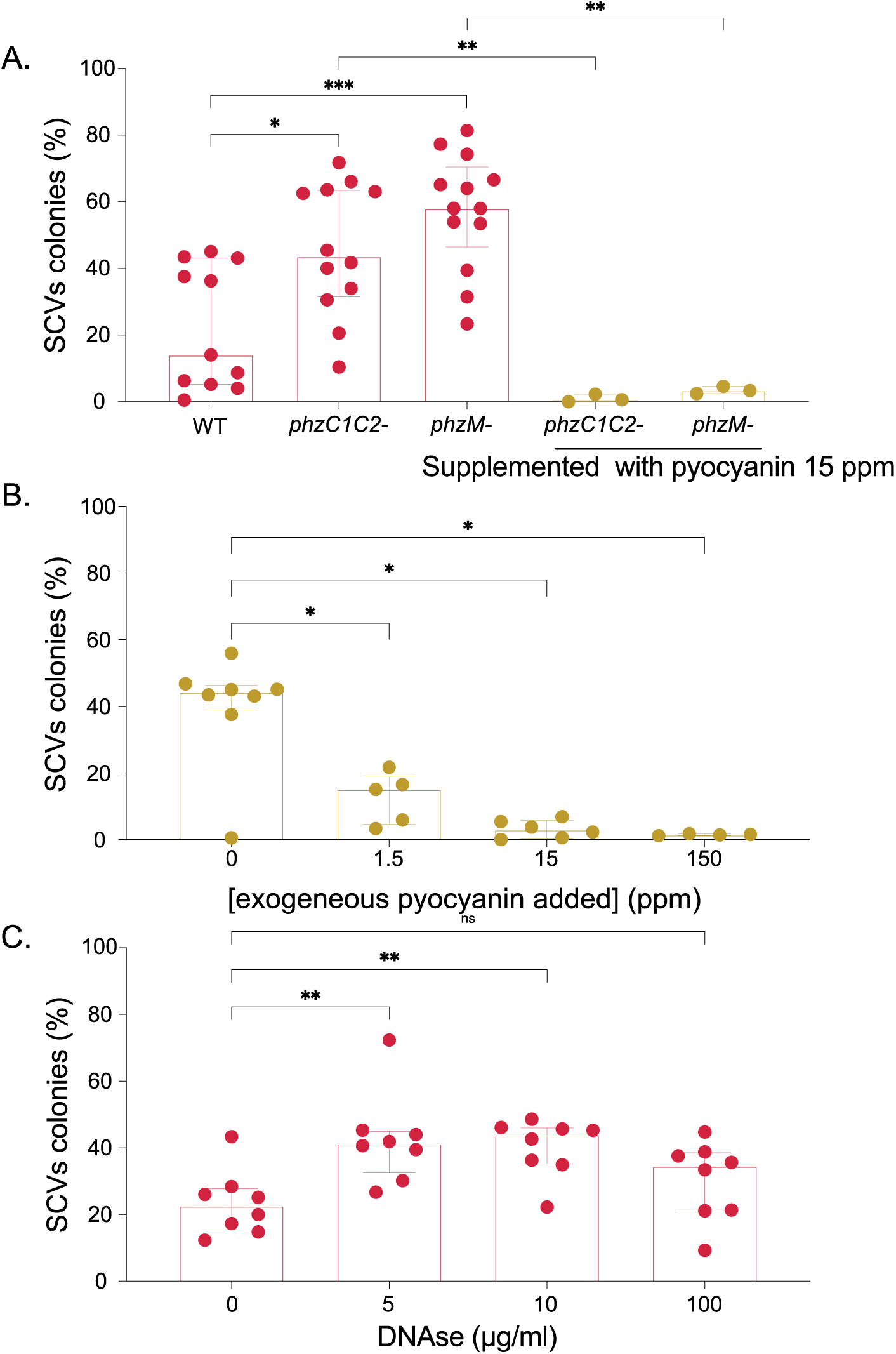
SCVs rate in cultures with an electron shuttle defect in PA14 phenazine mutants and pyocyanin or DNAse supplemented cultures. (A) PA14 WT, PA14 *phzC1C2*^*-*^ and PA14 *phzM*^*-*^ were inoculated in TSB (pink) or TSB supplemented with 15 ppm of purified pyocyanin (yellow). (B) PA14 WT was inoculated in TSB supplemented with 0, 1.5, 15 or 150 ppm of purified pyocyanin. (C) PA14 WT was inoculated in TSB with 0, 5, 10, 100 µg/mL of DNAse and incubated in static conditions. After 48 hours of incubation in static conditions, cultures were spread onto TSA plates and % of SCVs colonies in total population were quantified. Stars represent the statistical significance of the results calculated by a non-parametric Mann-Whitney test, ****, P Value ≤ 0.0001; ***, P Value ≤ 0.001; **, P Value ≤ 0.01; *, P Value ≤ 0.05; ns, not significant.

To determine if the absence of an efficient electron shuttle, rather than the absence of pyocyanin itself, was responsible for the higher SCVs rate, we next focused on extracellular DNA (eDNA), which enhances electron shuttle (35). Interestingly, addition of DNAse doubled the percentage of SCVs in standing cultures (Fig. 4C), indicating that reduction in eDNA, likely causing a reduced electron shuttle efficiency, promotes the emergence of SCVs. All together, these results indicate that the absence of an efficient electron shuttle to palliate limitations in O_2_ availability leads to increased SCVs emergence. Supporting this conclusion, we observed that SCV colony-biofilms displayed wrinkles at their surface, contrasting with the smooth surface of parental strain colony (Fig. 5).

**Fig. 5.**
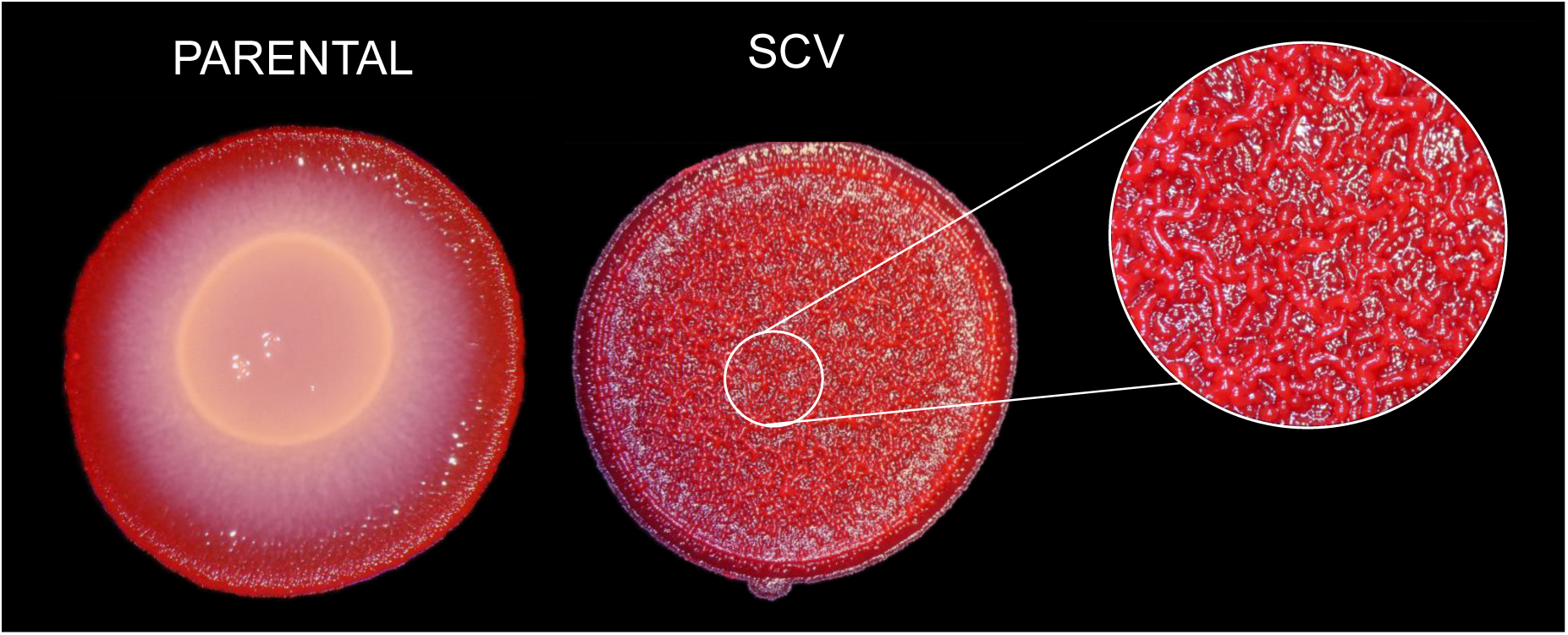
Morphology of PA14 parental and SCV colonies on Congo Red plates. Plates were observed with a binocular StemiDV4 (Zeiss) and photos were taken with a DMC-ZS60 camera (Panasonic Lumix), after 24 h of incubation at 30°C.

### Emergence of SCVs rebalances the redox state

Restrictions in electron acceptors unbalances the intracellular redox state. Pyocyanin facilitates redox balancing, particularly in the absence of electron acceptors. Since the lack of both O_2_ or pyocyanin promotes SCVs emergence, we hypothesized that the intracellular redox state could be directly correlated with SCVs emergence. Indeed, in standing cultures in which O_2_ availability was limited, the NADH/NAD^+^ ratio was 14 times higher during the first 15 h of incubation compared with a 24 h standard agitated culture in which electron acceptors are not limiting (Fig. 6A). Strikingly, the NADH/NAD^+^ ratio of the cell population was drastically reduced upon emergence of SCVs to finally reach the low NADH/NAD^+^ ratio of the agitated culture (Fig. 6A), indicating that selection for SCVs correlates with the intracellular oxidation state.

**Fig. 6.**
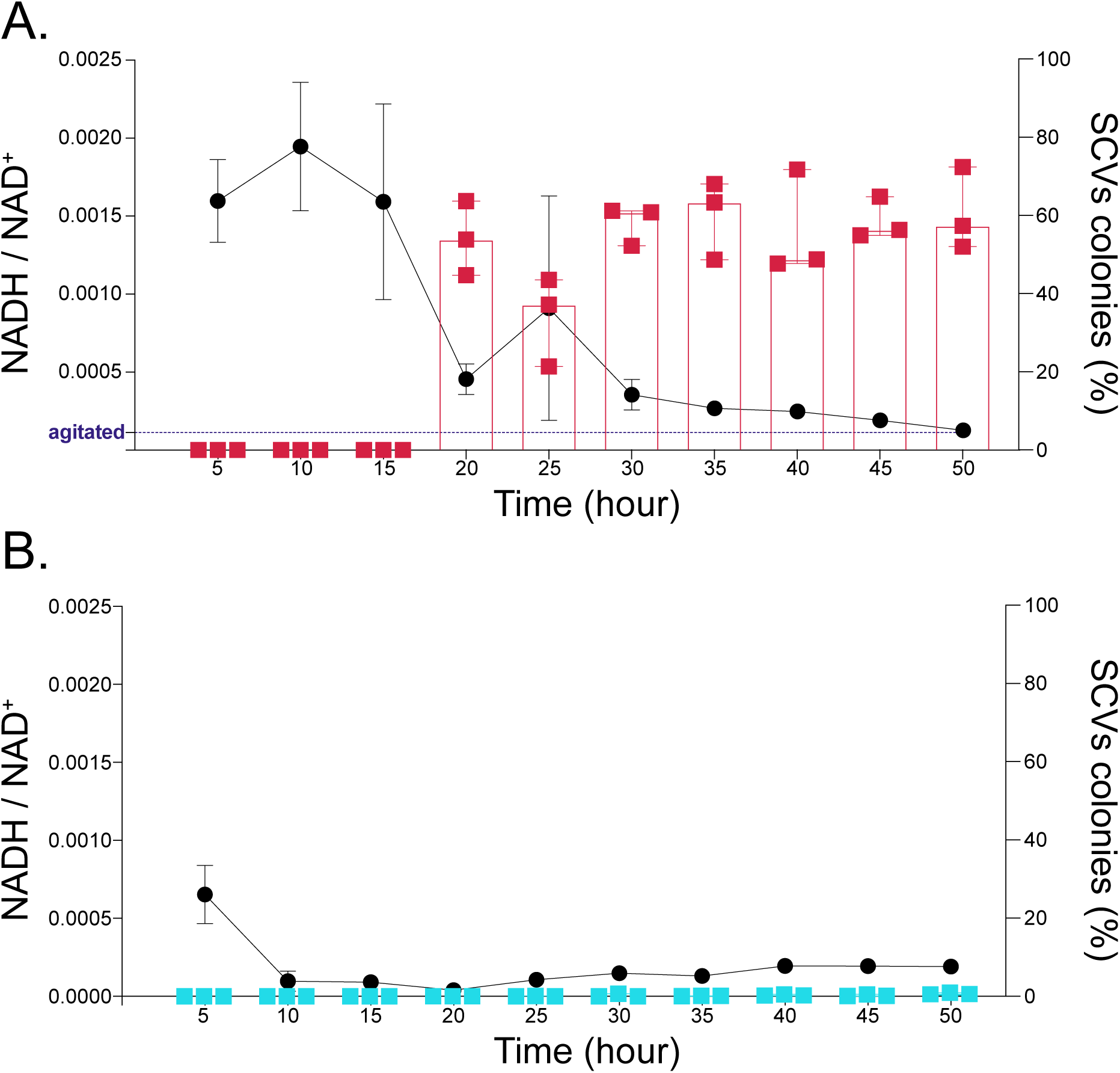
Effect of SCV emergence and KNO_3_ supplementation on the intracellular redox state of standing cultures. PA14 WT was inoculated in (A) TSB or (B) TSB supplemented with 50 mM KNO_3_. Cultures were incubated in static conditions for 50 h. Cultures were sampled each 5 hours and the % of SCVs colonies in total population was determined (■ symbols, histogram). Concomitantly, NADH and NAD^+^ were extracted from the static cultures to assess redox intracellular state (● symbols, curve). Purple dash line in (A) indicates the NADH / NAD^+^ ratio of an overnight PA14 culture incubated in agitated conditions.

The same experiment was performed in TSB supplemented with KNO_3_. As expected, the NADH/NAD^+^ ratio was only slightly increased early on during the exponential growth phase, but rapidly returned to the basal level corresponding to a 24 h standard agitated culture, confirming that the intracellular redox state is not unbalanced in conditions where the availability of electron acceptors is not limiting (Fig. 6B). Since SCVs emergence is significantly reduced in static cultures supplemented with KNO_3_ (Fig. 3 and Fig. 6B), an imbalance in intracellular redox state could represent the condition promoting the emergence of SCVs.

## DISCUSSION

Energy generation in *P. aeruginosa* is mainly based on oxidative substrate catabolism. In the respiratory cascade, O_2_ is the terminal electron acceptor (36). When O_2_ becomes limiting, intracellular accumulation of reduced compounds destabilizes the redox state, disturbing essential metabolic functions. Thus, maintaining a balanced cellular redox state is required for the survival of the cell. For instance, in surface growing biofilms, an O_2_ gradient is forming from the top to the bottom of dense microbial communities (37). Thus, bacteria can quickly exhaust available electron acceptors. To ensure its survival, *P. aeruginosa* has developed different strategies to palliate low-O_2_ conditions and rebalance redox intracellular state in biofilms when the use of an alternative electron acceptor is impossible, such as phenazines production (17, 18) and increase of the O_2-_ exposed surface (20, 21).

Here, we have cultivated *P. aeruginosa* PA14 in standing cultures, an *in vitro* mode of growth where O_2_ is limiting. We observed spontaneous emergence of SCVs in an O_2_-depleted culture, whereas it is abolished when an alternative electron acceptor is usable in presence of a functional Anr regulator. SCVs emergence was also correlated with a rebalance of the intracellular redox state. Since redox state imbalance is a direct consequence of O_2_ limitation, emergence of SCVs is likely to be an adaptive strategy to cope with poor O_2_ availability. Also, SCVs emergence rate was directly linked to the strength of the survival pressure induced by O_2_ limitation, as indicated by the increase in SCV proportion under microaerobic conditions or in absence of phenazines. Dietrich *et al*. previously concluded that availability of O_2_ and phenazine modulates colony morphology on agar, probably to increase the exposure surface and palliate O_2_ limitation (20). The surface of an SCV colony-biofilm is similar to the wrinkly colony described for phenazine-defective mutants or under low O_2_ conditions. Maybe the wrinkles could be the result of SCVs emergence in the colony?

Besides O_2_ limitation, the SCV phenotype can also be the results of exposures to different stresses. In one of the first report of SCV phenotype, Déziel *et al*. obtained SCVs by growing *P. aeruginosa* on an extremely hydrophobic source of carbon, hexadecane (31). In addition, accumulation of the toxic intermediate, gluconolactone, following knock out of the gluconolactonase coding gene *ppgL*, resulted in SCV emergence (38). Exposure to sub-lethal hydrogen peroxide level (H_2_O_2_) concentration during 120 generations also led to the emergence of SCVs (39). In all these scenarios, forming a biofilm is the best solution to ensure survival. For instance, to grow on hexadecane, the only way to thrive was to grow directly attached to the substrate, thus the need for rapid biofilm formation. This is similar to our O_2_ depleted standing cultures, in which growth as a biofilm at the air-liquid interface allows for increased access to O_2_. Thus, SCVs emergence could rather not only be a way to promote biofilm formation, but instead an essential process to readily form a biofilm and ensure survival, or a way to adapt to various stress conditions. Supporting this hypothesis, SCVs display a lot of specific properties that promote efficient biofilm formation compare to parental colonies (40): cell surface hyperpiliation and stronger adherence to abiotic surfaces (25, 31, 33) but also overproduction of exopolysaccharides (EPS) (Pel and Psl), the major component of the biofilm matrix, (27, 41).

In *Pseudomonas aeruginosa* natural habitats, SCV emergence could play a key role in adaptive strategy. One of the most studied *P. aeruginosa* habitats is CF infected hosts. *P. aeruginosa* is a key pathogen in CF individuals. Upon infection, *P. aeruginosa* colonizes CF mucus, a specific microenvironment displaying O_2_ gradient with O_2_-limited environments to anaerobic environments (42). *P. aeruginosa* can grow in the hypoxic/anaerobic CF mucus. Anaerobic growth may be in part supported by the terminal electron acceptor nitrate, allowing denitrification to generate energy (42, 43). Interestingly, SCVs have been reported several times from CF samples (25, 28–30) and could also be part of the redox stress response to hypoxia, largely responsible for the formation of biofilm, the predominant phenotype of *P. aeruginosa* in CF airways (44). Thus, we speculate that the stress of hypoxic/anaerobic environments induces *P. aeruginosa* to acquire phenotypic features, such as switch to the SCV phenotype, that allows it to enhance biofilm formation and evade hosts defenses and antimicrobial treatments, ensuring its survival. A correlation between the emergence of *P. aeruginosa* SCVs and infection persistence in animal models was established, supporting the idea that the SCV phenotype confers a fitness advantage under chronic infection conditions (45–47). Since *P. aeruginosa* is the main cause of morbidity and mortality in CF individuals and infections are becoming more difficult to treat because of the resistance to many antibiotics (48, 49), a better understanding of the pathways that contribute to survival and virulence of *P. aeruginosa* during infection, e.g. mechanisms of adaptation in O_2_-starved environments, is central to developing strategies to combat human *P. aeruginosa* infection.

In conclusion, our data show that *P. aeruginosa* PA14 SCVs emerge in response to an imbalanced redox state resulting from limitations in electron acceptor availability and that their emergence is correlated with formation of a surface pellicle, i.e. a biofilm-like structure, at the air-liquid interface. This allows access to O_2_ and a rebalance of the intracellular state, ensuring survival of *P. aeruginosa*. Here we pointed out a specific stressing condition leading to SCV emergence, but other stresses could result in SCV emergence. We propose that SCVs emergence is likely to be central in the adaptive response of *P. aeruginosa* accounting for its versatility and persistence in a variety of environments.

## MATERIALS AND METHODS

### Bacterial strains and liquid growth conditions

Bacterial strains are listed in Table 1. “Agitated culture” refers to strains grown in 5 ml tryptic soy broth (TSB; BD) at 37°C in a TC-7 roller drum (New Brunswick) at 240 rpm for 16 h. “Standing culture” refers to strains grown in static conditions at 34°C for up to 96 h. Standing cultures were grown in 5-or 8-ml TSB. When needed, TSB was supplemented with 50 mM KNO_3_, purified pyocyanin (chloroform extraction from a PA14 culture) or DNAse (Sigma). Standing cultures were inoculated at an initial OD_600_ of 0.05 and placed in an Infors incubator (Multitron Pro) without agitation, corresponding to “aerobic conditions”, or in an anaerobic jar with a lighted candle to consume ambient O_2_, corresponding to “microaerobic conditions”.

**Table 1.** Strains used in this study.

| **Strains *P. aeruginosa*** | **Lab collection**<br>**strain number** | **Reference** |
| --- | --- | --- |
| **PA14 Wild Type** | ED14 | (11) |
| **PA14 *anr::MrT7*** | ED1092 | (51) |
| **PA14 $\Delta\Delta phzC1C2$** | ED85 | Unpublished lab strain |
| **PA14 *phzM::Gm*** | ED120 | Gift from Pierre Cornelis |

### Quantification of SCVs

To increase reproducibility of the results, cultures were sampled at ~5 mm under the surface (underneath the pellicle), unless otherwise stated. A sterile P1000 tip with a cut end was added in the culture tube prior to inoculation to facilitate sampling through the tip and avoiding collection of the sticky pellicle. About 100-150 µl of the culture were sampled with a sterile Pasteur pipette and thoroughly mixed by pipetting up and down to homogenise sample and get rid of potential aggregates. Twenty µl were collected from the homogenised sample and diluted in TSB. Finally, 50 µl of the diluted sample were spread on tryptic soy agar plates solidified with 2% agar (TSA; AlphaBiosciences). Plates were incubated for 24 h at 30°C. CFU (Total + SCVs) colonies were counted manually from photographs using ImageJ. Results are expressed in SCVs percentage of total population. All experiments were performed at least in triplicates.

### Detection of O_2_ availability

To assess O_2_ consumption, we used the O_2_ sensitive dye resazurin. In presence of O_2_, blue resazurin is irreversibly reduced to pink colored resorufin, which is further reversibly reduced to colorless dihydroresorufin under reducing conditions, such as when O_2_ is depleted. TSB was supplemented with 100 mg/ml of sterile resazurin prior to inoculation of standing cultures. Resazurin reduction was assessed visually.

### Bradford protein assay

Due to the highly aggregative properties of SCVs, OD_600_ measurements were not appropriate to evaluate growth of standing cultures. The Bradford protein assay was used to quantify the total proteins concentration in all our samples. Pellets from 1 ml of culture were resuspended in 1 ml 0.1 N NaOH and incubated 1 h at 70°C, then protein concentrations were measured in samples according to the manufacturer guidelines for the Bradford reagent (Alfa Aesar).

### Extraction and quantification of intracellular of NADH and NAD^+^

Extraction of NADH and NAD^+^ was carried out according to the method described by Kern *et al*. (50). Briefly, 5 ml standing cultures were vortexed. Two 1.8 ml samples of culture were centrifugated 1 min at 16,000 x *g*. Pellets were resuspended in 200 µl 0.2 M NaOH (for extraction of NADH) or 200 µl of 0.2 M HCl (for extraction of NAD^+^). The extracts were incubated 10 min at 50°C, followed by 5 min on ice. To neutralize solutions, 200 µl 0.1M HCl or 200 µl 0.1M NaOH were added one droplet at the time while vortexing. Samples were centrifugated 5 min at 16,000 x *g* and the resulting supernatants were collected and stored at −80°C until quantification. For quantification, each step was performed protected from light. Eighty µl of a master reagent mix (20 µl bicine buffer (1M, pH 8, Sigma-Aldrich), 10 µl H_2_O, 10 µl EDTA 40 mM, 10 µl ethanol 100%, 10 µl thiazolyl blue tetrazolium bromide (Sigma-Aldrich) 4.2 mM and 20 µl phenazine ethosulfate (Sigma-Aldrich) 16.6 mM) were dispensed into individual wells of a 96-well plate. Fifteen µl of samples or standards (pure NADH and NAD^+^, Sigma-Aldrich) were added to each well. Plate was incubated 10 min at 30°C. Reaction was started by addition of 5 µl alcohol deshydrogenase (Sigma-Aldrich A-3263) (1 mg/ml in 0.1M bicine (pH 8)). The absorbance was measured at 570 nm every 30 sec during 45 min with a microplate reader (Cytation3, Biotek). Slopes were generated from standards and samples absorbance measures at 570 nm over time. Standard curves were generated from NADH and NAD^+^ standard slopes and used to calculate absolute concentrations of NADH and NAD^+^ in samples. Values were normalized to the protein concentration of the original cell culture sample and correlated with % SCVs in the same cell culture.

## ACKNOWLEDGMENTS

This work was supported by grant MOP-142466 from the Canadian Institutes of Health Research (CIHR). Dr. Alison Besse was a Fellow of the Calmette & Yersin postdoctoral grant from the Institut Pasteur International Network. The funders had no role in study design, data collection and interpretation, or the decision to submit the work for publication.

AB, MCG and ED conceived the project, contributed to experimental design and interpreted results. AB contributed to data acquisition. AB wrote the manuscript. MCG and ED reviewed and edited the manuscript.

